# DrTransformer: Heuristic cotranscriptional RNA folding using the nearest neighbor energy model

**DOI:** 10.1101/2022.09.08.507181

**Authors:** Stefan Badelt, Ronny Lorenz, Ivo L. Hofacker

## Abstract

**Background:** Folding during transcription can have an important influence on the structure and function of ℝNA molecules, as regions closer to the 5’ end can fold into metastable structures before potentially stronger interactions with the 3’ end become available. Thermodynamic ℝNA folding models are not suitable to analyze this problem, as they can only calculate properties of the equilibrium distribution. Other software packages that simulate the kinetic process of ℝNA folding during transcription exist, but they are mostly applicable for short sequences.

**Results:** We present a new algorithm that tracks changes to the ℝNA secondary structure ensemble during transcription. At every transcription step, new representative local minima are identified, a neighborhood relation is defined and transition rates are estimated for kinetic simulations. After every simulation, a part of the ensemble is removed and the remainder is used to search for new potentially relevant structures. The presented algorithm is deterministic (up to numeric instabilities of simulations), fast (in comparison with existing methods), and it is capable of folding ℝNAs much longer than 200 nucleotides.

**Availability:** This software is open-source and available at https://github.com/ViennaRNA/drtransformer.

## 1 Introduction

Most common RNA secondary structure prediction models calculate the thermodynamic minimum free energy (MFE) structure, which assumes that (a) the whole molecule is available, and (b) the molecule is given sufficient time to fold into the optimal structure. However, cells synthesize RNA molecules in 5’ to 3’ direction via transcription: the RNA polymerase reads the DNA template and appends single nucleotides to the RNA molecule at a rate between 20 and 200 nucleotides per second, although pausing during transcription can last for multiple seconds (Pan and Sosnick, 2006). Typically, the last 14-18 nucleotides are assumed to be caged from the polymerase and prevented from base-pairing, the remaining part of the RNA can fold freely before the full molecule is available.

The expected time for RNA structure formation ranges over many orders of magnitude: from fast hairpin formation and general helix zipping reactions on the order of microseconds (Pörschke, 1974; Ma *et al*., 2006), branch migration reactions on the order of milliseconds, to other complex rearrangements which may take on the order of seconds or much longer. Accordingly, many experimental findings show that the RNA structures forming during transcription can influence the conformation found at the end of transcription, for example: folding paths prevent the MFE structure formation (Kramer and Mills, 1981; Xayaphoummine *et al*., 2007), folding paths speed up MFE structure formation (Heilman-Miller and Woodson, 2003), pausing sites assist the folding large molecules (Wong *et al*., 2007), formation of hairpin structures cause the termination of transcription (Roberts, 2019).

In silico modeling of cotranscriptional folding is algorithmically challenging, as the ensemble of relevant structures at any particular transcription step can be overwhelming both in terms of computational and visual analysis. Even if only two structures dominate the ensemble in terms of occupancy, many more intermediate structures may have to be included in a model to estimate the dynamics between these two structures. Among existing algorithms are stochastic simulations (`Kinfold` (Flamm *et al*., 2000), `Kinefold` (Xayaphoummine *et al*., 2005), `RNAkinetics` (Danilova *et al*., 2006)), master equation methods (`BarMap` (Hofacker *et al*., 2010), theoretical work from (Zhao *et al*., 2011)), the deterministic prediction of a single folding trajectory (`Kinwalker` (Geis *et al*., 2008)), as well as a recent model to interpret experimental data `R2D2` (Yu *et al*., 2021) and theoretical work on combining stochastic modeling with deterministic helix kinetics (Xu *et al*., 2022).

The stochastic simulator `Kinfold` presents the simplest model where changes in secondary structure correspond to elementary moves, i.e. opening and closing of single base-pairs. The main difficulty with this approach, however, is that many simulations are needed to get statistically significant results, leading to an overwhelming amount of data without further post processing. Typically, both the generation and analysis of `Kinfold` data is time consuming and challenging for users.

Here, we present `DrTransformer`, short for “DNA-to-RNA Transformer”: a heuristic for cotranscriptional folding to provide fast approximations of `Kinfold` simulations. The software is open source and specifically designed with an easy usage interface to make cotranscriptional folding simulations more accessible to the community. We show that the results of `DrTransformer` compare well to statistically correct sampling of folding trajectories of short sequences. The accuracy of simulations, as well as the limits of sequence length in practice are heavily dependent on structural diversity, cotranscriptional folding traps, and on the chosen parameters. Our concluding runtime estimate uses natural group II intron RNA sequences of 620-781 nucleotides length to demonstrate applicablitiy far beyond the capabilities of any other competing methods besides the overly simplistic model `Kinwalker`.

## 2 Methods

Given a molecule of length *n*, the `DrTransformer` algorithm proceeds via *n* nucleotide extension cycles (see Fig. 1), which are composed of an *expansion step* where new candidate secondary structures are identified, their neighborhood relation is determined and transition rates are calculated, a *coarse graining step* where similar structures are combined into single representative local minimum conformations, a *deterministic simulation* to redistribute occupancies of secondary structures until the next nucleotide is transcribed, and a *pruning step* where structures with low occupancy are discarded. Naturally, the algorithm starts with expansion (from the first transcribed nucleotide) and ends with pruning (after the last simulation); we will use this order for discussing the procedures in detail, but start with some formal definitions and background information to level the ground for all sections to come.

**Figure 1:**
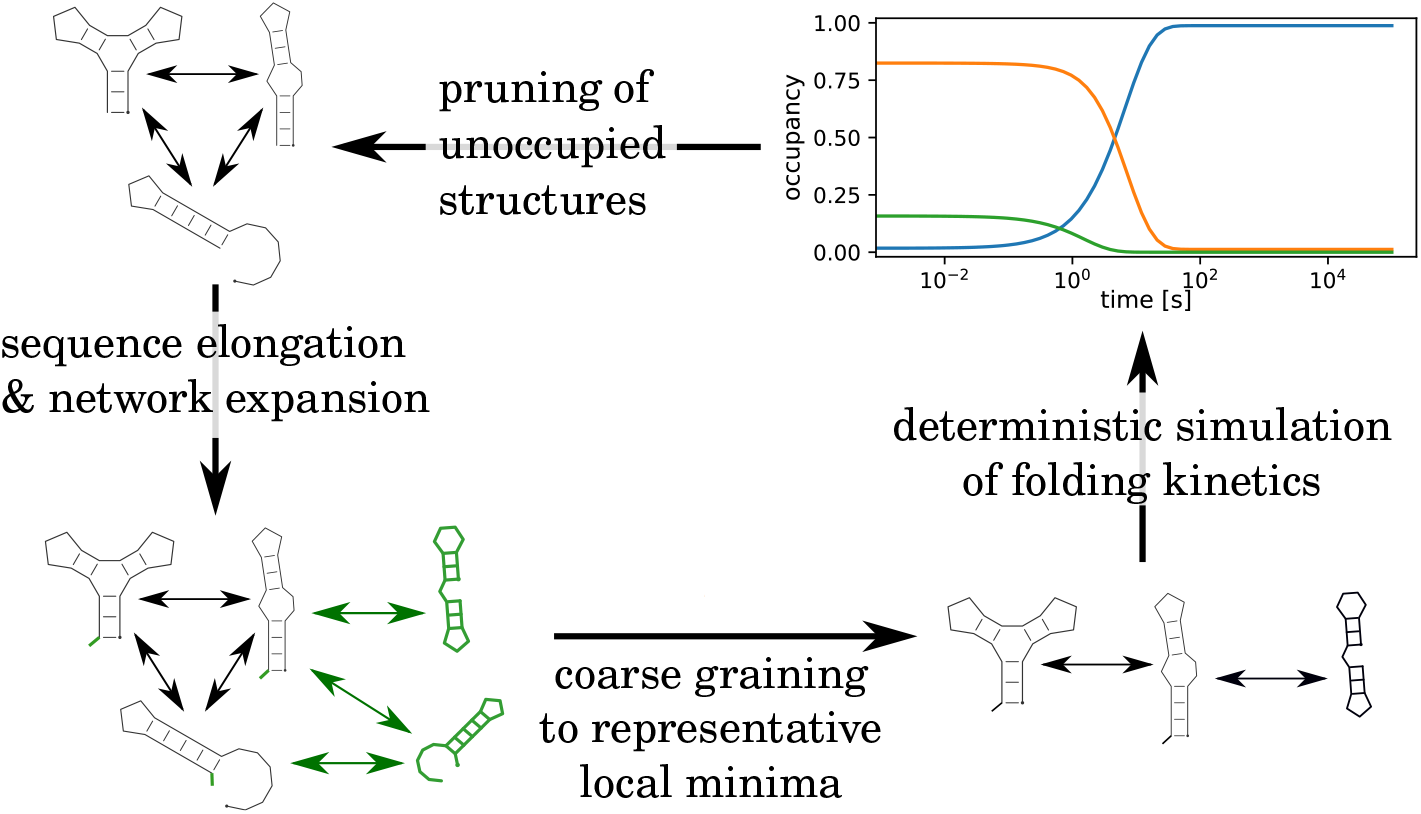
Bird-eyes view on the `DrTransformer` algorithm. For every new nucleotide, the sequence is elongated and the network of possible structures and transitions is expanded. A coarse graining procedure identifies local minimum conformations that are good representatives of the present network. A deterministic, kinetic simulation is used to determine how occupancies change in the present ensemble. Before the next nucleotide is transcribed, all unoccupied structures are removed from the network.

### 2.1 Background and Notation

We start with the notions of RNA sequence and structure for a molecule of length *n*.

#### Definition 1

*The* ***sequence*** *σ*_[1,*n*]_ *of an RNA molecule is an ordered list of n nucleotides from 5’ to 3’ end, where σ*_*i*_ ∈ {*A, C, G, U* }.

#### Definition 2

*The* ***structure*** *or* ***secondary structure*** *x corresponding to RNA sequence σ, is a set of base-pairs* (*i, j*), *subject to four conditions: (1) isosteric base-pairs only:* (*σ*_*i*_, *σ*_*j*_) ∈ {*(A,U), (U,A), (C,G), (G,C), (G,U), (U,G)* }, *(2) every base forms at most one pair: if* (*i, j*), (*i, k*) ∈ *x then j* = *k, (3) base-pairs have to be nested: if* (*i, j*) ∈ *and* (*p, q*) *and i < p < j then i < q < j, and (4) hairpins loops contain at least three unpaired nucleotides: if* (*i, j*) ∈ ∈ *x then* |*i − j*| *>* 3.

These definitions present a class of nucleic acid structures for which a thermodynamic energy model exists (Turner and Mathews, 2009), and enable the calculation of the MFE conformation as well as the partition function in *O*(*n*^3^) time and *O*(*n*^2^) space. `DrTransformer` uses the `ViennaRNA package` Lorenz *et al*. (2011) for secondary structure predictions. Notably, secondary structures as defined here do not include so-called pseudoknots (i.e. conformations containing non-nested base-pairs), and base-triplets (one base engaging in two pairs). Both motifs are present in many interesting RNA structures, but would require a more sophisticated energy evaluation model and substantial adaptations to the presented algorithm.

#### Definition 3

*An* ***energy landscape*** *ℒ* = (*𝒮, ℳ, E*) *is a directed, strongly connected graph, where the* ***nodes*** *x, y* ∈ *𝒮 are structures and* ***edges*** *m*_*x→y*_ ∈ *ℳ are “moves” corresponding to all possible* ***reactions*** *(i*.*e. direct transitions) between structures. m*_*x⇋y*_ *denotes a reversible pair of reactions. Every node x* ∈ *𝒮 has a fitness attribute in form of a thermodynamic* ***free energy*** *E*_*x*_.

Finding the “right” set of nodes, the “right” set of reactions, and the “right” rates for those reactions in combination with the “right” energy function is the main challenge for this work, and there exist many different approaches to find satisfying and/or practical definitions for those terms in the context of RNA energy landscapes, e.g. Flamm *et al*. (2000, 2002); Kucharik *et al*. (2014); Entzian *et al*. (2021). We follow the common assumption that the most accurate energy landscape model corresponds to *elementary* base-pair opening and closing steps. This is implemented in `Kinfold`, a Gillespie-type stochastic simulators for RNA folding, which infers rate constants from the Metropolis (Metropolis *et al*., 1953) or Kawasaki (Kawasaki, 1966) model. In later sections, we will discuss under which circumstances multiple elementary reactions can be combined into one overall reaction and then use the Arrhenius model to derive reaction rates for transitions that involve multiple base-pair opening and closing steps. The following definitions will be important for identifying repesentative structures (*x, y* ∈ *𝒮*), valid transition reactions between those structures (*m*_*x⇋y*_ ∈ *ℳ*) and reaction rate constants that are consistent with the energies from the thermodynamic nearest neighbor energy parameters.

#### Definition 4

*The* ***base-pair distance*** *d*(*x, y*) *between two structures x, y is the cardinality of the symmetric difference between the sets of base-pairs formed by x and y respectively*.

#### Definition 5

*An* ***elementary path*** *or P*_*xy*_ *is a sequence of distinct secondary structures, starting in x and ending in y which can be generated via single base-pair opening and closing steps. An elementary path of length m* = *d*(*x, y*) *is called* ***shortest path*** *or* ***direct path***, *otherwise it is an* ***indirect path***.

#### Definition 6

*The* ***saddle energy*** *ℰ*_*x⇋y*_ *of a path between two structures P*_*x⇋y*_*is is the maximum free energy on a path:* 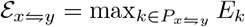

*A structure x is a* ***local minimum***, *if all immediate neighbors have equal or higher free energy. A structure x is a* ***δ-minimum*** *if there exists no structue y with E*_*y*_ *< E*_*x*_ *reachable by a path with saddle energy ℰ*_*x⇋y*_ *< δ*.

#### Definition 8

*The* ***occupancy*** *of a structure 𝒪*_*x*_ ∈ ℝ^[0,1]^ *is a real-valued probability of observing the structure*.

Since the occupancy is a probability, 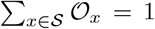. We use the term occupancy, to avoid confusion with the thermodynamic equilibrium probability of a structure.

### 2.2 Expansion algorithm

At the start of transcription, before any structures are occupied, the expansion routine yields only the current minimum free energy structure (typically one unpaired nucleotide). At subsequent iterations, every currently occupied structure gets an unpaired nucleotide attached which turns it into a so-called parent conformation^1^, and then the expansion algorithm proceeds via multiple stages to build a representative landscape for simulation: Fist, a heuristic identifies new, presumably relevant, secondary structures given the current parent conformations. Second, base-pair distances between all occupied and new structures determine a “guiding neighborhood” relation and constrained folding (Lorenz *et al*., 2016) is used to identifying additional relevant structures. Third, saddle energies, and local minima are identified along direct paths for all transitions in the guiding neighborhood; the neighborhood relations are refined to account for additional local minima and reaction rates are calculated for all transitions with suitable elementary path profiles.

#### 2.2.1 Secondary structure search

The core of the secondary structure search procedure is based on the observation that (in the nearest neighbor model) a newly transcribed nucleotide can only interact with bases in the exterior loop, i.e. all nucleotides of the RNA molecule that are not already enclosed by a base-pair, otherwise a forbidden pseudoknotted structure would be formed. Hence, the structure search is focused on local conformation changes around the exterior loop which are triggered by the newly transcribed base.

Our procedure for generating new candidate structures from a parent conformation uses constrained MFE folding and assumes that only a few base pairs of the parent conformation can be opened within the time availale for one elongation step. We thus translate the parent structure into multiple constraints that keep all base pairs and loop regions constant, except for the exterior loop and (different combinations of) helices adjacent to the exterior loop (see Fig. 2). If MFE folding using these constraints yields a new (energetically equivalent or better) structure, then it is returned by the structure search algorithm. (A parameter determines the minimum number of freed bases during helix fraying. If less bases are freed, then any nested helix is considered to fray as well.) Additionally, the procedure returns the (unconstrained) MFE structure for the current sequence length. Note that we can reduce the runtime by removing enclosed (constant) helices during constrained folding. This reduces the sequence length *n* to the length of the exterior loop, which is effectively constant and independent of the overall transcript length.

**Figure 2:**
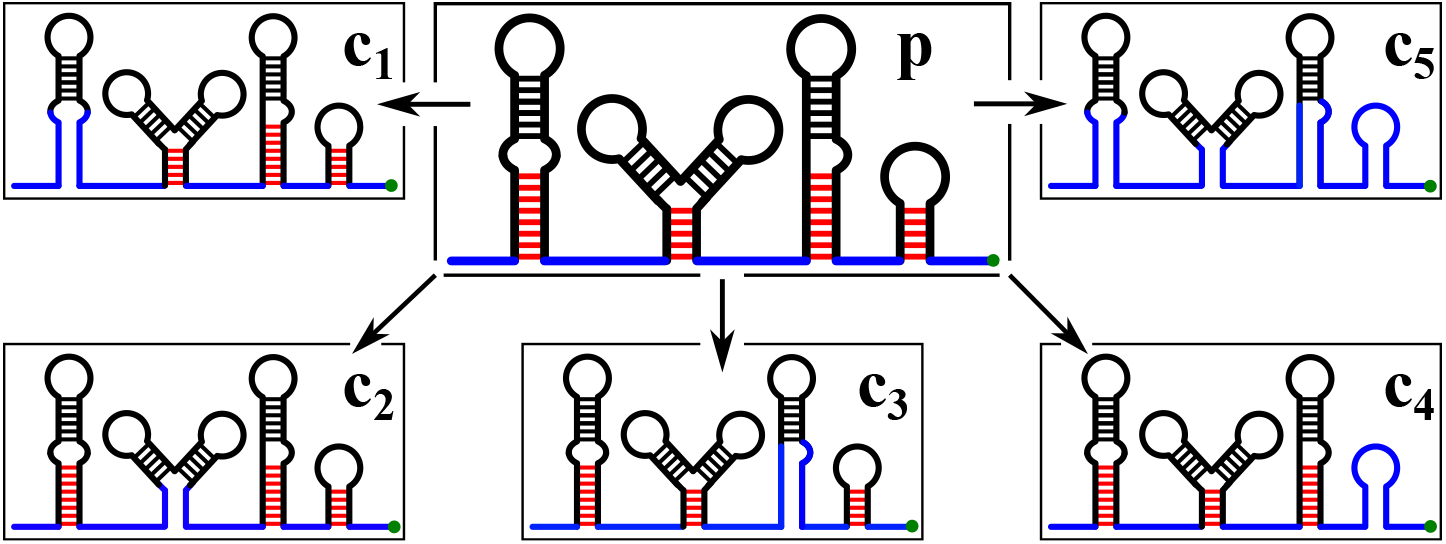
Constraints to generate new relevant secondary structures. The parent structure *p* has a new unpaired base attached (green dot) which can pair with any available base in the exterior loop (blue). Fraying helices are shown with red base-pairs, black base-pairs are part of enclosed helices. Each fraying helix is opened separately to produce constraints *c*_1_, *c*_2_, *c*_3_, *c*_4_, and all fraying helices are opened at once to produce constraint *c*_5_. The latter allows for rearrangements involving two (or more) competing fraying helices.

#### 2.2.2 Guiding neighborhood construction

Guiding neighborhood construction is the first step for finding a neighborhood relation between all deemed relevant structures, and it is independent of known transitions from previous iterations of the algorithm. The approach is iterative and proceeds in three steps. First, the set of currently relevant structures is extended through constrained MFE folding: every structure is used as a constraint to find potentially better structures which are compatible with the constraint. Second, base-pair distances are used to find a neighborhood relation. Formally, a pair of reversible guide edges *g*_*x⇋y*_ is added to the landscape, if there exists no structure *i*, such that

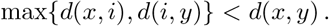

Subsequently, for every node *i*, for every pair of guide edge neighbors *x, y*, new shortcut edges *s*_*x⇋y*_ are added if

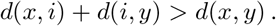

Third, for every guide and shortcut edge, the MFE structure along the direct path is calculated. This is done by constrained folding using only the union of base-pairs of start and end structures, all other base-pairs are forbidden. If the MFE structure along the direct path does not equal starting or end structure, it is included into the set of relevant structures. If new structures are found in this third step, then the whole procedure is repeated, otherwise the algorithm terminates.

##### Intuitions behind guide and shortcut edges

Intuitively, all structures are connected with guide edges, *unless* there exists a structure *in between*. This approach guarantees that every structure is reachable by every other structure in the landscape, and, in the limit of the full set of secondary structures (Wuchty *et al*., 1999), this approach yields the elementary base-pair opening and closing move set used by statistically accurate stochastic simulators of RNA folding (Flamm *et al*., 2000; Schaeffer *et al*., 2015). However, the approach has drawbacks if most structures are not explicitly included. As an example, consider the structures `x=..((…))`, `y=..(….).`, and `z=(((…)))`; even though an elementary path for a transistion *m*_*xz*_ does not have to visit species *y*, only the guide edges *g*_*x⇋y*_ and *g*_*y⇋z*_ are part of the network. Connecting “neighbors of neighbors” whenever there exists a shorter path is a comparatively simple solution that fixes the problem stated above, while maintaining correctness in the limit of the full set of suboptimal structures where no shortcut edges would be added.

#### 2.2.3 Estimating transition rates between structures

The last part of network expansion is to analyze folding paths and estimate reasonable transition rates. It is worth emphasizing that the following algorithm assumes that all important representative structures are already known, but additional intermediate structuctures are included whenever the probability of a transition cannot be described by a single rate constant. We provide an approximate model based on biophysical intuitions which may lay the foundations for a more rigorous analysis that is beyond the scope of this contribution. Metaphorically, the structure search procedure reveals some “islands of interest” in a vast “ocean” of secondary structures, and the guide edges give us a smart guess on which islands are closest to each other. Our task – as ambitious cartographers – is to set sail and record as much information as possible about (i) which islands can be reached *directly* (i.e. without visiting other islands in between) and (ii) what is the success probability of reaching specific island *y* after setting sail from *x*. It is obvious that every piece of collected information gets us closer to the ground truth system, and that any information collected in the past (at a previous transcription step) is still valid in any future transcription step.

For every guide and shortcut edge, a path *P*_*x⇋y*_ is generated using the `findpath` heuristic (Flamm *et al*., 2001); `findpath` searches for a path with minimal saddle energy among all direct paths. The path presents a one-dimensional energy landscape where *δ*-minima and saddle energies (see Sec. 2.1) can be determined using a flooding algorithm (see Fig. 3). Only if the path has no *δ*-minimum *k ≠ x ≠ y* with *E*_*k*_ *≤* max(*E*_*x*_, *E*_*y*_), a direct transition has been found. In that case, two valid transition edges *m*_*x⇋y*_ are added to the set of edges ℳ and the saddle energy ℰ_*x⇋y*_ is used to calculate a reaction rate constant using the Arrhenius model

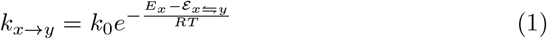

where the pre-exponential factor *k*_0_ is a rate constant to map simulated time scales to the wall-clock time observed in experiments, *R* is the gas constant and *T* is the temperature. Otherwise, if new *δ*-minima *k ≠ x ≠ y* with *E*_*k*_*≤* max(*E*_*x*_, *E*_*y*_) are found, then those structures are included into the model and new saddle energies are calculated for the subpaths. (Only the saddle energy for the overall path has been optimized by the `findpath` algorithm, which is why some subpaths must be evaluated again to find a lower saddle energy.)

**Figure 3:**
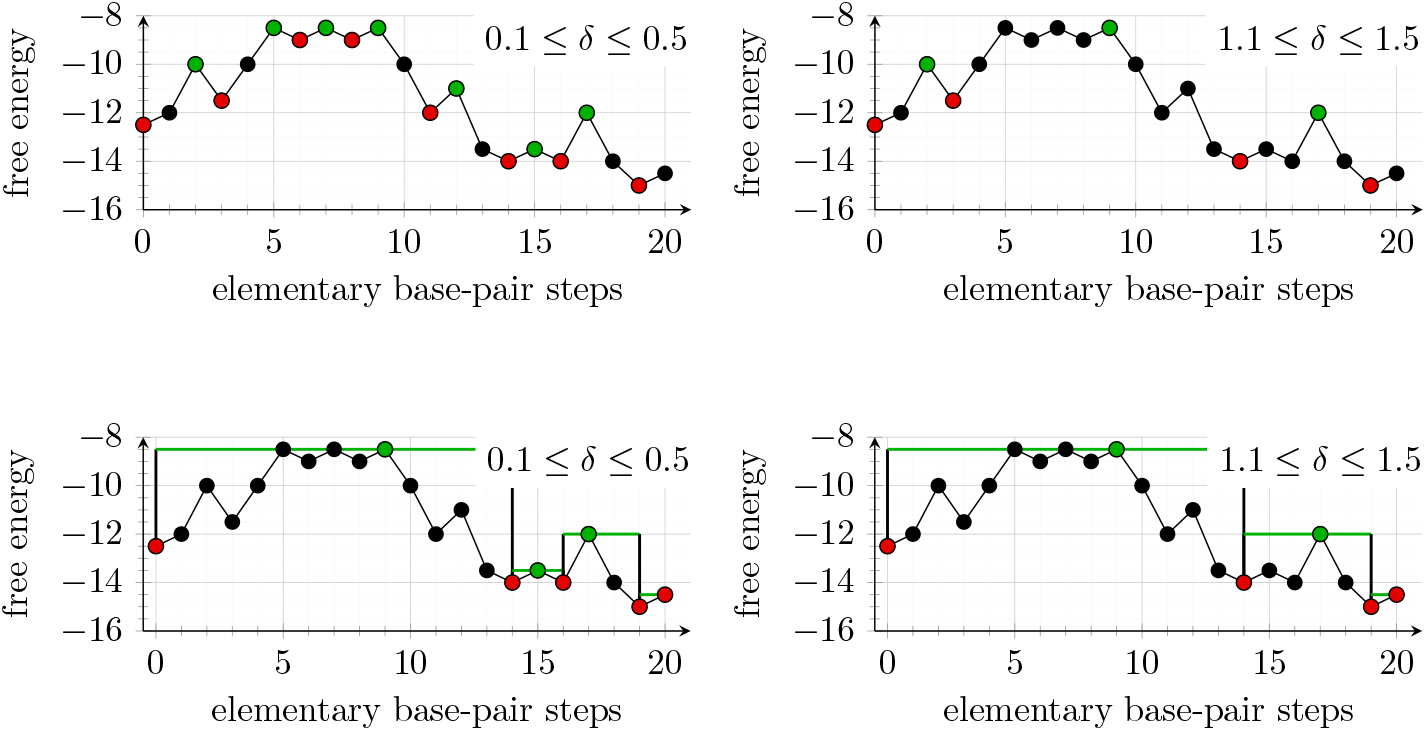
Results of the path flooding procedure and the corresponding model of direct transitions between *δ*-minima. All images show the free energy change along a direct path with 20 elementary steps. Every point corresponds to a structure, *δ*-minima are highlighted in red and saddle points are highlighted in green. **(top row)** *δ*-minima and saddle points as identified by the flooding procedure with 0.1 ≤ *δ* ≤ 0.5 and 1.1 ≤ *δ* ≤ 1.5, respectively. The coloring assumes that nodes at the same energy level have processed from left to right by the flooding procedure. **(bottom row)** Direct transitions included in the expanded network for the given *δ*. All *δ*-minima with energy lower than starting and end conformation are explicitly included as relevant secondary structures, the saddle energy (green line) is used to estimate transition rates between *δ*-minima.

The model uses the minimal observed saddle energy on a valid path to calculate (or update) the rate. At this stage, start and end structures *x, y* always remain explicit nodes in the landscape, even if flooding reveals that they are not *δ*-minima. Such cases will be dealt with later in the coarse graining procedure, as there might be multiple competing *δ*-minima to which a structure can get assigned to.

Note that we can store the saddle energies corresponding to a path for future calculations, because a path will never become invalid. In other words, if there exists an edge *m*_*x⇋y*_ in the graph, there must exist a path *P*_*x⇋y*_ that does not pass through a *δ*-minimum *k* with *E*_*k*_ *≤* max(*E*_*x*_, *E*_*y*_). If a guide edge *g*_*xy*_ has been found and the transition edge between parent structures *m*_*x⇋y*_ is known from an earlier transcription step, then there is no need to calculate it again. Also, a previous parent transition *m*_*x⇋y*_ remains part of the landscape even if the edge cannot be found in the guide graph.

##### Intuitions behind the transition rate model

In combination with the guide graph construction and in the limit of the full suboptimal structure space, this rate model yields the elementary base-pair move model with Metropolis rates. However, the model is applied in a sparse structure space, where elementary paths determine the transition rate. If a *δ*-minimum *k* has energy *E*_*k*_ *>* max(*E*_*x*_, *E*_*y*_), then it is assumed to be short-lived, and the overall reaction rate between structures *x* and *y* is dominated by the saddle energy ℰ_*x⇋y*_. Otherwise, if *k* has energy *E*_*k*_ ≤ max(*E*_*x*_, *E*_*y*_), then this assumption cannot be made and structure *k* has to be included as species into the model.

It is worth pointing out that the condition *E*_*k*_ ≤ max(*E*_*x*_, *E*_*y*_) should be taken with reservations. It is not guaranteed that a *δ*-minimum *k* must be short-lived, just because it happens to be on an elementary path between lower energy *δ*-minima. The crutial distinction for our purposes is that there are some cases in which *k* is already an explicit species in the model, e.g. because it is sufficiently occupied from a former transcription step, and other cases where the only evidence that *k* may be of interest is that it has been discovered on an elementary path *m*_*x⇋y*_. The latter is not per-se an interesting structure and it may not yield a better rate estimate for transition *m*_*x⇋y*_ than the saddle energy ℰ_*x⇋y*_.

### 2.3 Coarse graining algorithm

Coarse graining reduces the model from many detailed structures to fewer abstract structures while preserving certain quantities of interest. This section describes an algorithm to (i) reduce the (potentially large) number of secondary structures found during expansion to a smaller set of representative conformations, (ii) approximate the dynamics between the remaining relevant conformations assuming elementary base-pair transitions. Thus, the algorithm serves two purposes: a reduction of the model and the elimination of bias in the set of representative structures introduced during the expansion process.

The enabling concept is a timescale separation. Reactions between two structures that are separated by a low energy barrier are assumed to be fast (effectively instantaneous), reactions between all other structures have to be modeled explicitly (see Eq. 1 for rate calculation). Eventually, coarse graining yields a set of *δ*-minimum structures which should be understood as abstract species, (i.e. representatives of sets of detailed structures); all other structures are transient structures, which are not explicitly modeled, but they are used to determine the transition rates between *δ*-minimum structures.

#### 2.3.1 Top-down coarse graining

Fig. 4 illustrates the algorithm using a simple one-dimensional toy energy landscape, the generalization to high dimensions will be described below. Note that this is the same landscape as in Fig. 3, where we describe a flooding algorithm to identify *δ*-minima on direct paths. The top-down coarse graining algorithm processes a list of structures sorted from high energy to low energy (the order of structures at the same energy level is irrelevant). For every conformation *k*, take the set of outgoing reactions that yield a conformation with lower or equal energy. If any of these reactions has saddle energy ℰ_*x⇋y*_ *< E*_*k*_ + *δ* (i.e. it is *fast*), then *k* is a transient structure which will be removed from the model, otherwise it is a *δ*-minimum and the algorithm continues to process the next structure *k*.

**Figure 4:**
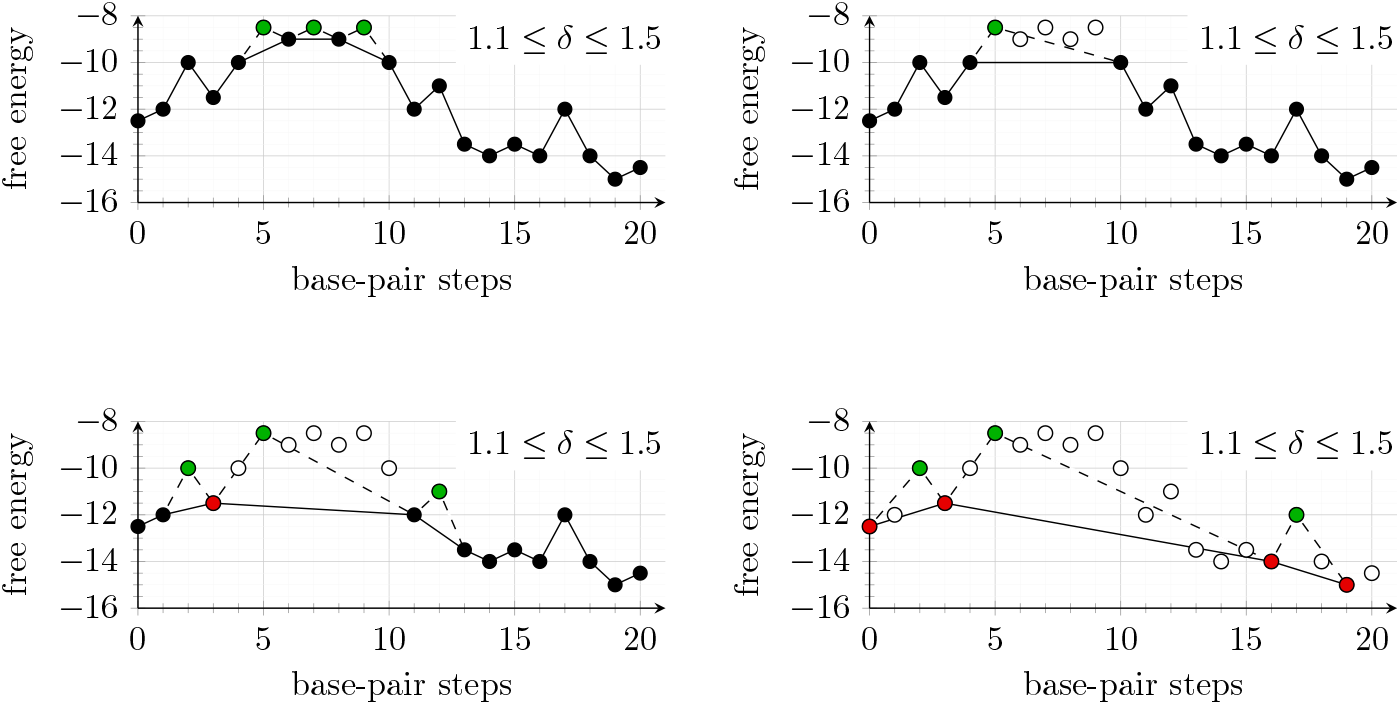
The top-down coarse graining procedure illustrated using an hypothetical path example. Four characteristic steps are shown. Top left, the three structures with highest energy have been processed. They are removed from the set of structures, but their energy still determines the transition rate between neighbors. Top right, two more structures have been removed, only one transition is left and the energy barrier is given by a green saddle point structure. Bottom left, the first *δ*-minimum has been identified (red). This structure cannot be removed from the system because it has no outgoing fast reaction. Bottom right, the final coarse graining consists of four *δ*-minimum conformations and three reversible direct transitions where the rate is calculated from the energy of saddle points (green).

If *k* is a transient structure, then its occupancy is divided among the neighbors *x* where ℰ_*x⇋y*_ is minimal. Note that all higher energy neighbors must be *δ*-minima, otherwise they would have been removed from the landscape (a *δ*-minimum cannot be reached by a fast reaction from a structure with lower energy). Every neighbor *x* of *k* which is reachable by a fast reaction is connected with all other neighbors *y* of *k*. In other words, if two neighbors *x, y* are both only reachable by slow reactions, then they are not connected.

#### Limitations of top-down coarse graining

Notably, this model does not account for the entropy of abstract structures, i.e. the partition function of all structures represented by *δ*-minima. Adapting the model to incorporate entropy is not straight-forward, as both the free energies of *δ*-minima as well as transition rates have to be adjusted accordingly.

### 2.4 Kinetic simulation

The coarse grained network of *δ*-minima and corresponding transition reactions is written into a rate matrix

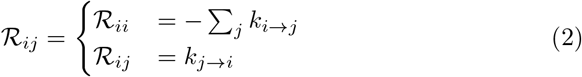

where every row *i* corresponds to the constants of the linear equation 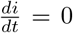 that must be satisfied at thermodynamic equilibrium *t → ∞*. In combination with the vector of initial occupancies 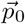, any vector 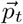 can be derived using

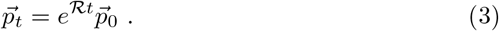

where the matrix exponential *e*^*ℛt*^ can be calculated efficiently in various ways for less than order 10,000 species (Moler and Van Loan, 2003). The two specific approches described below have been implemented previously in the program `treekin` (Wolfinger *et al*., 2004), but `DrTransformer` provides a standalone implementation of this process using the well-known Python libraries `numpy` (Har-ris *et al*., 2020) and `scipy` (Virtanen *et al*., 2020). First, *e*^*ℛt*^ can be calculated directly using the Pade approximation which is remarkably stable against numeric instabilities, but comparatively slow when calculating many different time points. The more efficient solution decomposes the matrix into a matrix of eigenvectors *S* and a diagonal matrix of eigenvalues Λ to solve the equation *e*^*ℛt*^ = *Se*^Λ*t*^*S*^*−*1^. In this solution, matrix decomposition is the time consuming part, but *e*^Λ*t*^ for every time-point *t* can be calculated in linear time. In order to counteract numeric instabilities and avoid complex solutions, we first use the detailed balance property of our system (*P*_*i*_*k*_*ij*_ = *P*_*j*_*k*_*ji*_) to derive the equilibrium distribution vector 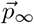, and to derive a symmetric matrix matrix *U* = Ω^*−*1^*ℛ*Ω, where Ω is a diagonal matrix with elements 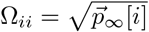. Now the symmetric matrix is decomposed into eigenvectors and eigenvalues *U* = *S*Λ*S*^*−*1^, and Eq. 3 can then be rewritten as

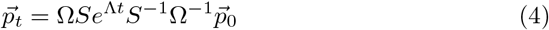

where Ω*S* and 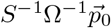 are constant with respect to changes over time *t*.

Every simulation has a linear regime [*t*_0_, *t*_1_] and a logarithmic regime (*t*_1_, *t*_8_]. The linear regime simulates folding kinetics for the time inteval of nucleotide extension, e.g. *t*_1_ = 0.02 seconds at a transcription rate of 50 nucleotides per second. During transcription, the logarithmic regime is used to “look ahead” until the end of transcription, i.e. 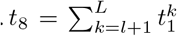 where *l* is the length of the current trancript, *L* is the length of the full sequence, and 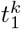 is *t*_1_ at length *k*. This look-ahead simulation is used to find structures which should be exempt from pruning (which will be discribed in Sec. 2.5), because they reach a relevant occupancy at the time scale of transcription. After transcription, *t*_8_ is set to the final post-transcriptional simulation time.

### 2.5 Pruning algorithm

Pruning connects simulation results (Sec. 2.4) with the next round of graph expansion (Sec. 2.2) and the algorithm is partly intertwined with of both of those processes. The general idea is that occupancies of individual secondary structures change, and low-populated structures are discarded in order to keep the system computationally tractable.

The core procedure is remarkably simple, all prunable structures are sorted from lowest to highest occupancy, and marked for deletion as long as their combined occupancy remains below a threshold ∑𝒪_*x*_ ≤ *o*. When structures are deleted, their occupancy is distributed evenly among all remaining neighboring structures, or recursively, among neighbors of neighbors if all of the neighboring structures are also marked for deletion.

Whether a structure is prunable, depends on whether its occupancy remained under that same theshold *o* during the look-ahead simulation described in Sec. 2.4. Whether a structure that is marked for deletion is actually deleted, depends on whether it is revisited during the next graph expansion procedure. Thus, the pruning procedure does not delete structures but labels them as inactive, which ensures that previously collected data on transition rates, etc. remains cached. Only the structures that remain inactive during landscape expansion have their occuancy distributed among neighboring conformations. We provide an option to set the parameter *o* that specifies the maximum amount of density that can be discarded from the current occupancy distribution, and a parameter to remove structures from the cache, if they remain inactive for more than a certain number of extension steps.

## 3 Results

Cotranscriptional folding can have a strong influence on the predicted structure even for comparatively short sequences. The data from Fig. 5 suggests that less than 50% of random sequences with 120 nucleotides are at equilibrium 60 seconds after transcription.

**Figure 5:**
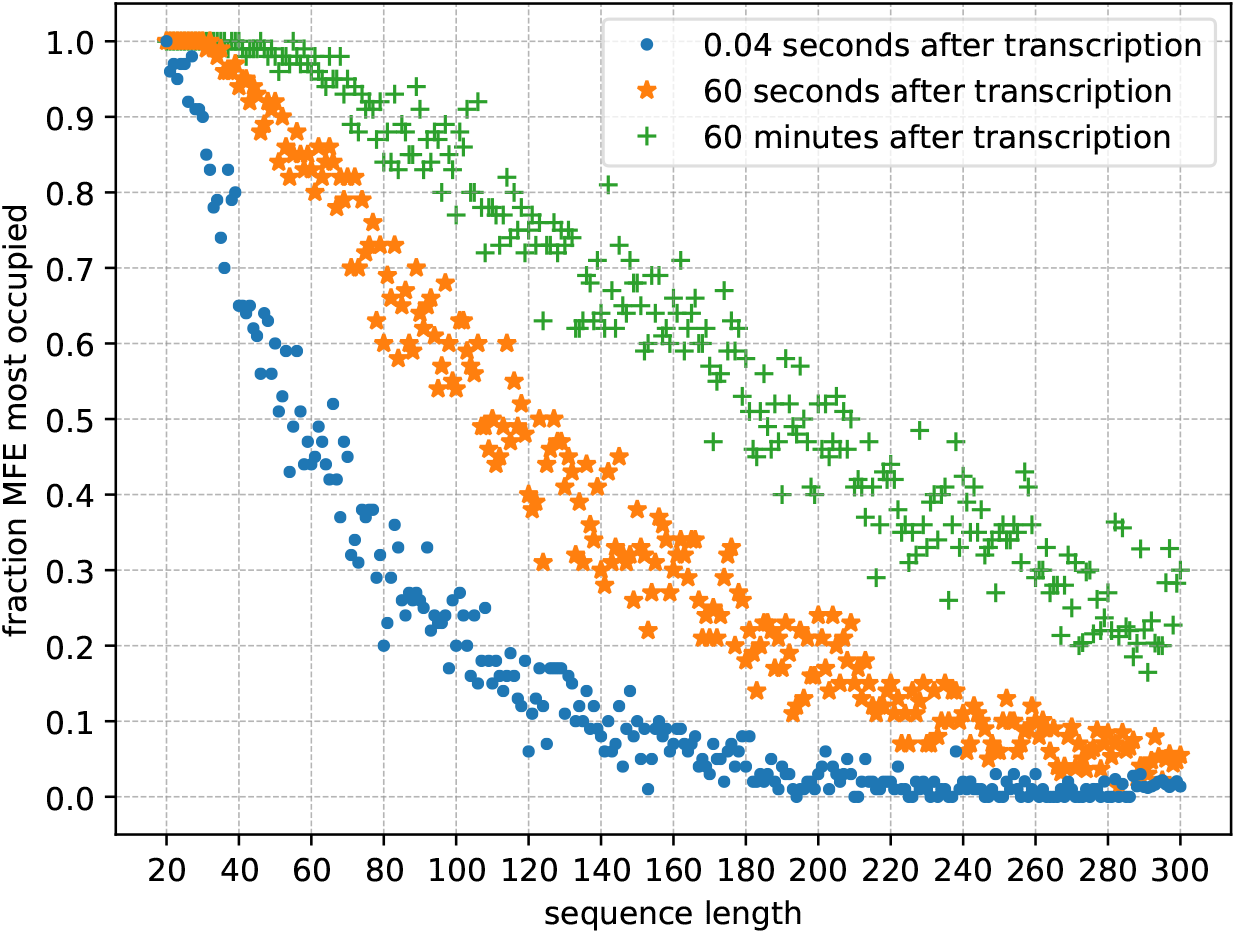
How often is the most occupied structure after transcription the MFE structure? We compare 0.04 seconds after transcription (the time at which the next nucleotide would be attached at a transcription rate of 25 nt/s), 60 seconds after transcription and 60 minutes after transcription. `DrTransformer` simulations suggest a rapid decline, e.g. less than 50% of 60 nt sequences are in their MFE conformation at the end of transcription, and less than 50% of 200 nt sequences are in their MFE conformation one hour after transcription. Each data point is the average of `DrTransformer` simulations for 100 random sequences of that length.

Generally, many parameters can influence `DrTransformer` simulations in subtle ways to drastically change the conformations found at the end of transcription. For example, the coarse graining strength reduces the number of explicitly modelled structures, the occupancy cutoff parameter may discard conformations before they become relevant for expanding the network. Obviously, the kinetic simulation time per nucleotide can have a particularly strong influence on cotranscriptional simulation results.

As `DrTransformer` is a heuristic approach to cotranscriptional folding on an energy landscape with elementary base-pair transitions, we focus on the differences of cotranscriptional ensemble predictions when comparing `DrTransformer` with the underlying ground truth model implemented in the program `Kinfold`. While we illustrate the differences using an experimentally verified system here, an additional analysis using random sequences assesses the diversity of cotranscriptional ensembles from `DrTransformer` and `Kinfold` and their correspondence to the equilibrium distribution. This analysis can be found in Suppl. Sec. 1.

### 3.1 Estimation of a suitable *k*_0_ parameter

For didactic purposes, we distinguish two parameters that influence the simulation time per nucleotide: The *k*_0_ rate constant of the Arrhenius-type model used by `DrTransformer` (see Eq. 1) translates arbitrary time units given by free energy differences to wall clock time in seconds, and the extension time specifies how many seconds to simulate per nucleotide. (It is obvious that those parameters are dependent, e.g. doubling both parameters yields the same simulation time per nucleotide, but it is more natural to fix *k*_0_ to a commonly accepted value, and then vary the transcription rate in terms of nucleotides per second.)

In past contributions analyzing cotranscriptional folding using `Kinfold` (Helmling *et al*., 2017) and `BarMap` (Badelt *et al*., 2015), 4000 arbitrary simulation time units per nucleotide were used in combination with the Metropolis rate model. As `DrTransformer` uses the same energy model and is an approximation to `Kinfold` simulations, we expect that the simulation time per nucleotide should be on the same order of magintude. In particluar, we expect that `DrTransformer` can produce similar results to `Kinfold` and that differences in simulation results may be compensated by minor adaptations to the *k*_0_ parameter.

The simulations in Fig. 6 correspond to three different RNA molecules from an experimental study (Xayaphoummine *et al*., 2007) that illustrates how helix competitions determine the structure formed at the end of transcription. Briefly, two sequences are composed of the same palindromic sub-sequences (A, B, C, D) in forward and reverse order (“ABCD” and “DCBA”); the third sequence (“DCMA”) has a point mutation which changes B to M. The experiment demonstrates how the order of helix formation determines which structures are formed at the end of transcription, an effect that cannot be observed with a thermodynamic equilibrium prediction.

**Figure 6:**
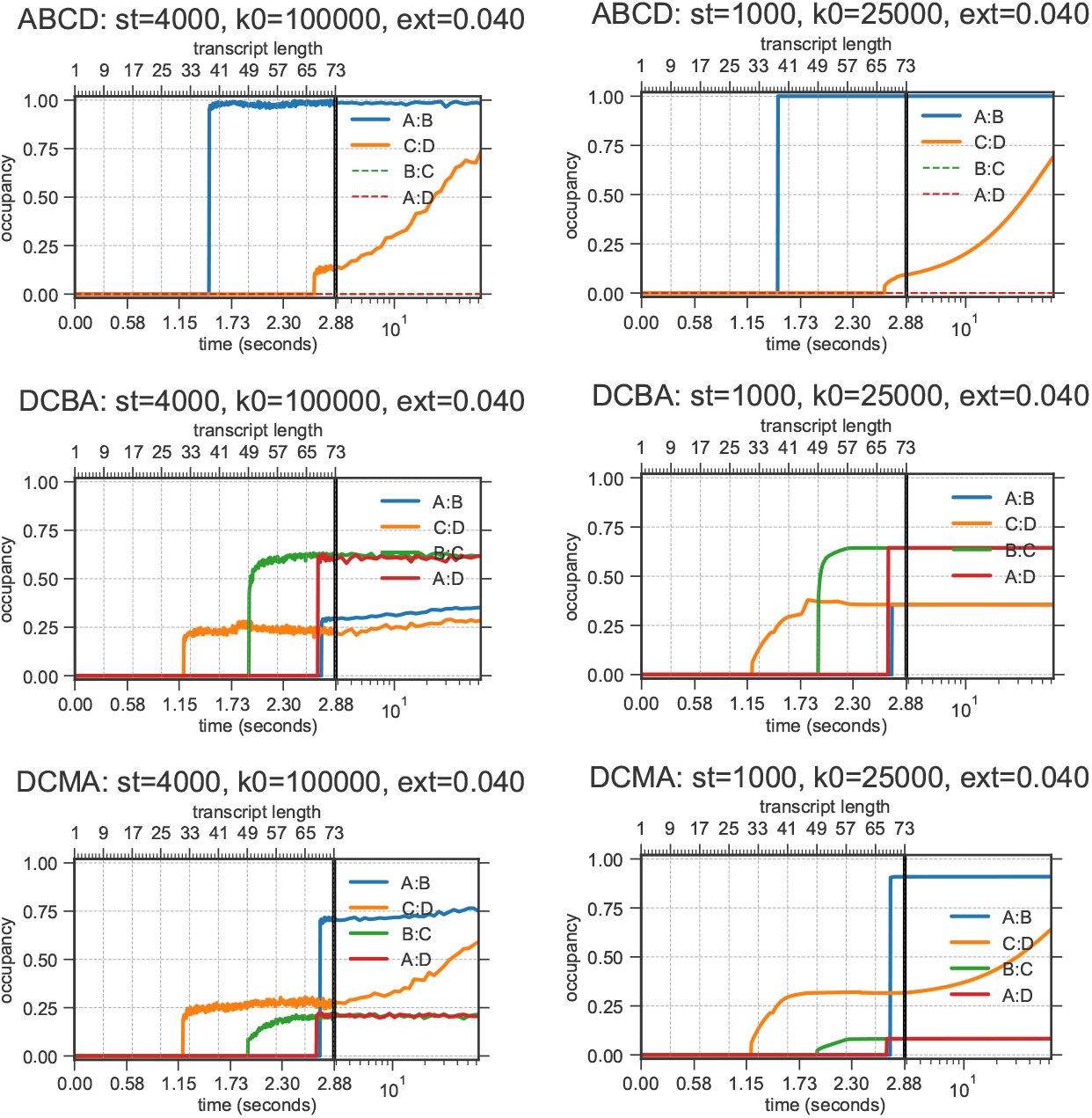
An example for adjusting the rate parameter *k*_0_ (which relates free energy differences to wall-clock time) to improve correspondence between `DrTransformer` and data. Here, we aim to match “ground truth” `Kinfold` simulations using sequences ‘ABCD’, ‘DCBA’, ‘DCMA’ from Xayaphoummine *et al*. (2007). (Note that this is much more difficult than fitting to the available experimental data for the structure distribution at the end of transcription.) The trajectories show fractions of structures that form helices “A:B”, “C:D”, “B:C”, and “A:D”. All simulations use exension time **ext**=0.040 s/nt, which corresonds to a transcription rate of 25 nt/s. **(first column)** `Kinfold` simulations using *k*_0_ = 10^5^. **(second column)** `DrTransformer` simulations using *k*_0_ = 2.5 · 10^4^. Varying *k*_0_ effectively changes the simulation time in arbitrary units per nucleotide (termed **st** in plot headers.) For further variation of simulation parameters see Suppl. Sec. 2.

For `Kinfold` simulations, we use the Metropolis rate model with parameter *k*_0_ = 10^5^ /s, in combination with a transcription rate of 25 n/s as suggested by Helmling *et al*. (2017). The `Kinfold` simulations explain experimental findings regarding the ensemble at the end of transcription (see Fig. 6, first column): ABCD folds almost exclusively into the MFE structure which forms only helices A:B and C:D, while the reverse sequence DCBA is cotranscriptionally trapped to form predominantly helices A:D and B:C. (Note that A:B is the same helix as B:A due to the palindromic sequences.) The single-base-mutation in DCMA decreases the effect of the cotranscriptional folding trap and helices A:B and C:D are again favored at the end of transcription. Fig. 6, second column, shows the cotranscriptional folding simulations of all three sequences using `DrTransformer`. Varying the simulation time per nucleotide (here by changing *k*_0_) yields a range of different simulation results. The simulations using *k*_0_ = 2.5 · 10^4^ /s come close to `Kinfold` predictions, suggesting that `DrTransformer` simulations for this example are approximately a factor 4 faster than `Kinfold` simulations. We show a additional variation of simulation parameters in Suppl. Fig. 1, 2, 3, 4, with a discussion on how they relate to experimental findings. (Interestingly, the ratio of structures forming *A*:*B* and *C*:*D* vs structures forming *A*:*D, C*:*B* in the *DCBA* molecule is heavily dependent on the transcription rate, an effect that can be observed from `DrTransformer` as well as `Kinfold` simulations, but – to our knowledge – has not been investigated experimentally.)

### 3.2 Performance analysis of DrTransformer

Fig. 7 shows runtime of `DrTransformer` as a function of sequence length. We compare two different datasets:

**Figure 7:**
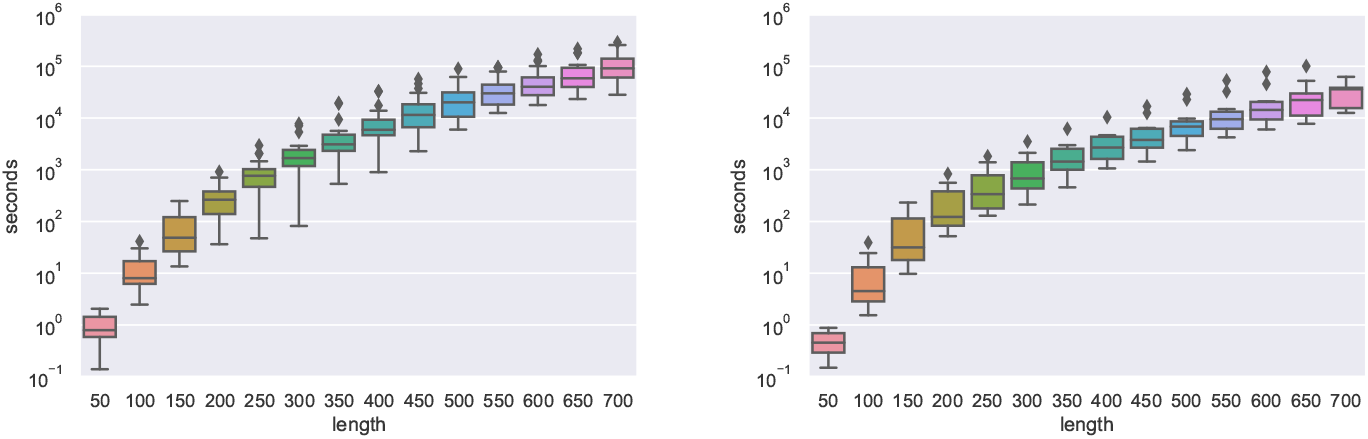
Runtime comparison of `DrTransformer` for random sequences up to 700 nt length and natural group II intron sequences (620 to 781 nucleotides length). Relevant non-default transcription parameters: `--o-prune` = 0.1.

- 20 random sequences up to a length of 700 nucleotides
- 11 group II intron sequences (620 to 781 nucleotides length)

The runtime can vary by more than an order of magnitude for different sequences of the same length. This is expected, as runtime depends on properties of the energy landscape, which are not known prior to the simulation. Also, the runtime for random sequences appears worse than for natural group II intron sequences. This may suggest that random sequences have a more diverse energy landscape and thus take longer to simulate. To produce the random sequence data plot, random 900 nt sequences were generated and given up to 7 days (≈6 *\** 10^5^ s) on a single core for `DrTransformer` simulations, respectively. Two out of 20 simulations did not terminate in time, one of them reached 896 nt length, the other only 743 nt. Hence, plots show the runtime up to 700 nucleotides. All group II intron sequences were simulated until the last nucleotide.

## 4 Discussion

We have presented a heuristic model for cotranscriptional folding that is applicable to both short and long RNA molecules. The Arrhenius-type model used by `DrTransformer` (see Eq. 1) is a generalized formulation of the Metropolis model, as it yields the same rates for single base-pair moves, but also allows for an estimation of transition rates that involve multiple base-pair insertions/deletions at once. While this model compresses elementary paths into single steps (which can speed up simulations) there may exist saddle structures with lower energy than those found on direct paths, in which case `DrTransformer` simulations would be slower than the corresponding `Kinfold` model. However, we have shown that, in practice, the model is able to qualitatively reproduce simulation results of the much more computationally expensive ground truth model `Kinfold` via slight adjustments to the *k*_0_ parameter.

### Usage notes

As mentioned for the sequences in Fig. 6, folding behavior can vary stronly dependent on the *k*_0_ parameter and the transcription rate – the latter is often not precisely known. Thus, it is recommended to vary the time per nucleotide to observe different types of structures at the end of transcription. If experimental results are known, the user can adjust the *k*_0_ parameter to plot results with a matching transcription rate. `DrTransformer` also provides options for pause sites at specific nucleotides, which in principle could be used to see how stochastic variations of the transcription rate (at each nucleotide) influence folding. Apart from options that influence the `DrTransformer` heuristic directly, `DrTransformer` provides an interface to all relevant `ViennaRNA package` energy model parameters such as temperature and alternative nearest neighbor parameters.

### Future work

The presumed ground truth Metropolis model for RNA folding is limited, as it depends on a single parameter to adjust how differences in free energy correspond to wall-clock time. It is also not clear whether intermediate steps during a helix-zipping reaction automatically correspond to well defined Markovian minima as presumed by the `Kinfold` model. `DrTransformer` provides a powerful open source toolbox to test new rate models in combination with experimental data on longer RNA molecules. For example, Zolaktaf *et al*. (2017) use an Arrhenius-type model for DNA folding kinetics with additional parameters for secondary structure context.

While this contribution focuses on the folding of single RNA molecules, we are planning to include the modeling of intermolecular nucleic acid interactions (small RNA influence on the cotranscriptional folding), and other ligand interactions. Incorporation of bimolecular interactions comes with subtle challenges concerning expansion, coarse graining and network pruning, but – as a first step – can follow a similar strategy as proposed by Wolfinger *et al*. (2018) who assume constant concentrations of the interacting molecule.

While `DrTransformer` provides parameters for simulations of long RNA molecules, more work is needed to determine parameters for which such predictions match experimental results. Many longer RNA molecules use specific types of pseudoknots to assist the folding into the target structure. Cotranscriptional folding of long RNAs may be especially interesting in the field of RNA origami (Geary *et al*., 2014), which typically relies on such types of interactions. While it seems difficult to improve `DrTransformer` predictions by allowing pseudoknots in general, a stepwise inclusion of certain classes in combination with a well described kinetic model is definitely an exciting direction.

## 5 Conclusion

`DrTransformer` presents a probabilistic model for cotranscriptional folding using heuristic energy landscapes for every transcription step. The program can be viewed as a hybrid approach between `BarMap`, a deterministic simulation on a-priori coarse-grained landscapes (Hofacker *et al*., 2010), and `Kinwalker`, a greedy algorithm to get the most probable trajectory (Geis *et al*., 2008). In practice `DrTransformer` can produce similar results to the most accurate model `Kinfold` (Flamm *et al*., 2000) using much less computation time, which allows users to scan over multiple possible transcription parameters quickly.

Immediate use cases for `DrTransformer` are the identification of cotranscriptional folding traps, and the analysis of cotranscriptional pausing sites with respect to secondary structure formation. More subtle analyses could involve the identification of sequence regions where certain transcription parameters are essential for correct folding, or where experimentally observed folds cannot be realized. The latter would suggest crucial interactions with unknown molecules that assist for correct folding. For example, it has recently been found that transient, non-native structures kick-start ribosome assembly (reviewed in Rodgers and Woodson (2021)). Finally, `DrTransformer` opens a variety of possibilities for sequence design, as the evaluation on whether an intended folding path is cotranscriptionally favorable is now much faster than using previous methods. For example, `DrTransformer` may be able to identify (and avoid) cotranscriptionally formed transcription termination motifs, which would greatly assist the design of large RNA molecules.

## Acknowledgements

We thank Christoph Flamm, Andrea Tanzer, and Michael T. Wolfinger for fruitful discussions in the early stages of the development of `DrTransformer`. SB wants to thank Erik Winfree for inspiring discussions on kinetic models for nucleic acid folding.

## Footnotes

1 Readers familiar with the nearest neighbor energy model will note that adding an unpaired base to the end of a structure can change its free energy due to so-called dangling end contributions. As a consequence, also energy barriers involving transitions of parent conformations may change. In order to save computation time, we introduce a small inaccuracy by evaluating the energy of any new structure at the full transcript length, assuming a tail of unpaired nucleotides whenever the transcript is shorter than the full length molecule.

