## Supplemental Material for "DrTransformer: Heuristic cotranscriptional RNA folding using the nearest neighbor energy model"

September 8, 2022

[1]Department of Theoretical Chemistry, University of Vienna, Austria.
[2]Division of Biology and Biological Engineering, California Institute of Technology, Pasadena, CA, USA
[3]Research Group Bioinformatics and Computational Biology, Faculty of Computer Science, University of Vienna, Austria

*To whom correspondence should be addressed

### 1 On the diversity of cotranscriptional ensembles

The following analysis uses an artificial dataset of 380 random sequences with lengths from 20 to 200 nucleotides. All sequences were simulated with `DrTransformer` and compared to the results of 200 stochastic cotranscriptional `Kinfold` simulations. First, we show how *diverse* ensembles are at the end of transcription compared to the ensemble at thermodynamic equilibrium. Ensemble diversity is calculated as the mean base-pair distance between randomly selected structures in the ensemble

$$\mathrm{MED}(A, B) = \sum_{a \in A} \sum_{b \in B} P_a P_b d(a, b) \tag{1}$$

were $P_a$ is the probability of structure $a$ (in ensemble $A$) and $d(a, b)$ is the base-pair distance. This formula can also be used to estimate the diversity within a single ensemble ($A = B$), by assuming that structures can be chosen multiple times with the same probability.

Equilibrium ensembles of random sequences are calculated using the `ViennaRNA package` and compared to the ensembles at the end of transcription using `Kinfold` and `DrTransformer`. The results are shown in Fig. 1. Under 60 nt length, random sequences are predominantly at equilibrium at the end of transcription, but for longer sequences the `Kinfold` ensemble after transcription is more diverse than the equilibrium distribution. Presumably, ensemble diversity is increased because sequences have not been designed to fold into a specific metastable structure and we observe an ensemble that contains some structures that are dominant at equilibrium, as well as others which are still trapped due to the history of the transcription process.

Perhaps surprisingly, one can see that length is a good predictor for ensemble diversity at end of cotranscriptional `Kinfold` simulations. This suggests that the main source of diversity comes from small variations at the level of individual base-pairs, which are expected to increase linearly with sequence length. `DrTransformer` groups similar structures into the same $\delta$-minimum, which means base-pair level variations are not observed. Consequently, length is not a good predictor of ensemble diversity. It appears as if the overall distribution of ensemble distances is similar for `DrTransformer` and equilibrium, but that structures with high diversity at the end of transcription can have low diversity at equilibrium and vice versa. In the direct comparison of cotranscriptional ensembles from `Kinfold` and `DrTransformer` we see that the `Kinfold` ensemble is more diverse (as expected), but we also now observe a trend where molecules with less diversity in `Kinfold` have less diversity in `DrTransformer`.

Finally, one can see that both `Kinfold` and `DrTransformer` ensembles at the end of transcription have a similar mean ensemble distance to the equilibrium ensemble. Presumably, base-pair level variations cancel when comparing `Kinfold` with equilibrium distributions, and the overall distances to dominant metastable structures dominate this analysis. This would suggest that `Kinfold` and `DrTransformer` are able to identify similar metastable structure candidates.

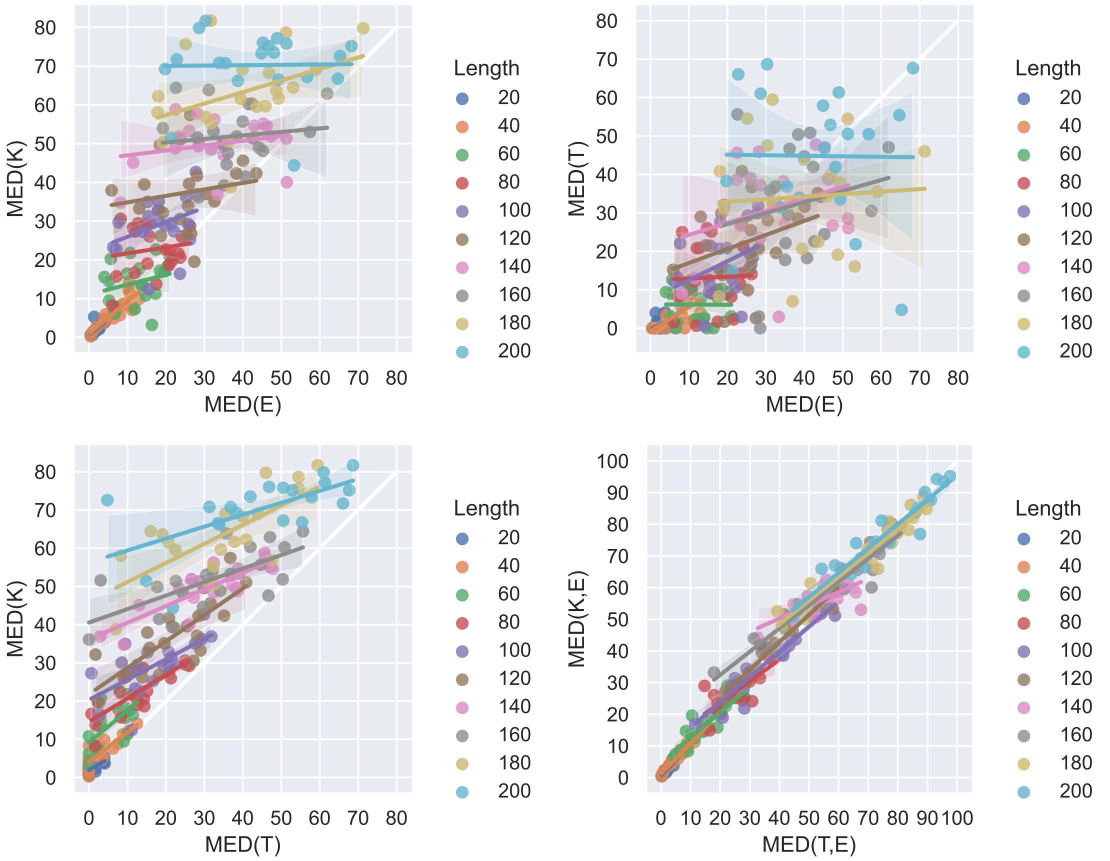


Figure 1: **Mean ensemble distances (MED).** Top left: The mean ensemble distance at the end of transcription calculated by `Kinfold` is larger than the mean ensemble distance at equilibrium. Within a specific length cohort, the ensemble distance at the end of transcription remains largely constant. Top right: The mean ensemble distance at the end of transcription calculated from `DrTransformer` simulations is distributed similar to the ensemble distance at equilibrium, but structures with high diversity at the end of transcription can have low diversity at equilibrium and vice versa. Bottom left: The mean ensemble distance at transcription end calculated by `Kinfold` and `DrTransformer` shows that `Kinfold` has a higher diversity, but it also shows that molecules with less diversity in `Kinfold` have less diversity in `DrTransformer`. Bottom right: The mean ensemble distance between `Kinfold` transcription end and equilibrium in comparison to the mean ensemble distance between `DrTransformer` transcription end and equilibrium. Both the `Kinfold` and the `DrTransformer` ensembles have similar distance to the equilibrium distribution.

#### 2 Comparison of `Kinfold` and `DrTransformer` by varying $k_0$

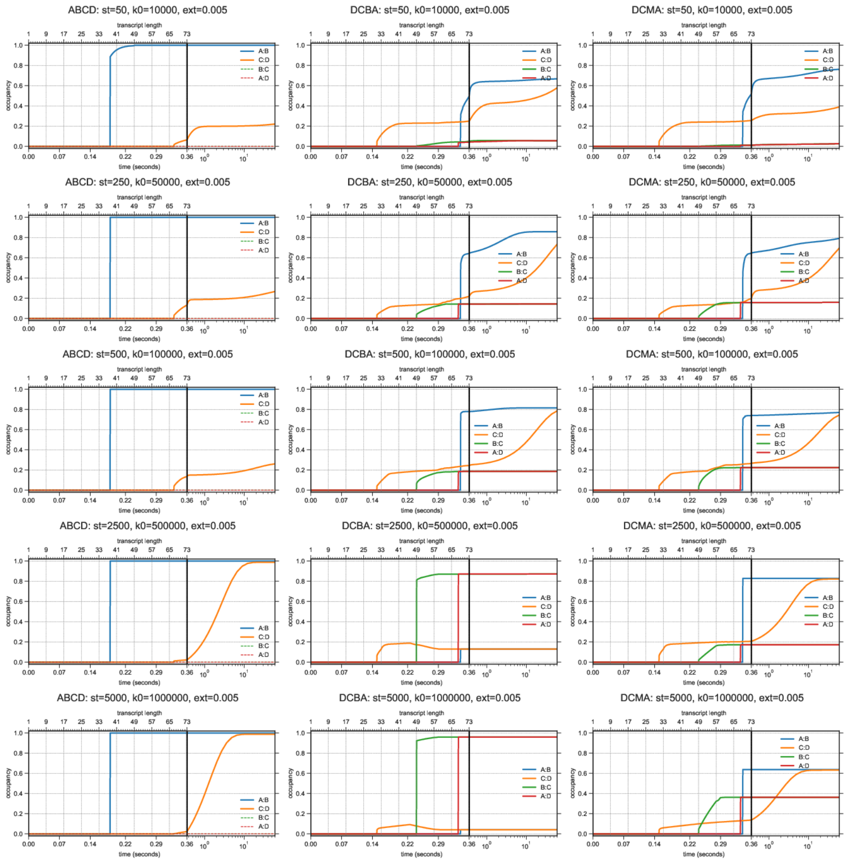


Figure 2: `DrTransformer` simulations of the three molecules 'ABCD', 'DCBA', 'DCMA' designed in **?** with varying simulation time per nucleotide. The extension time per nucleotides is constant at **ext**=0.005 s/nt which corresonds to a transcription rate of 200 nt/s, which is unusually high, but was suggested in the original publication. $k_0$ is varied by two orders of magnitude: **k0**=$\{10^4, 5 \cdot 10^4, 10^5, 5 \cdot 10^5, 10^6\}$, where $10^5$ is the `DrTransformer` default parameter. The simulation time per nucleotide in arbitrary units **st** is provided for comparison to a different transcrption rate in Suppl. Fig. 4. The corrsponding `Kinfold` simulations are shown in Suppl. Fig. 3. Experimental results for DCBA suggest 90% in B:C, A:D conformation and 10% in A:B, C:D conformation (**?**), which is in between the results for $k_0 = 5 \cdot 10^5$ and $k_0 = 10^6$. Accordingly, experimental results for DCMA suggest 50% in B:C, A:D conformation and 50% in A:B, C:D conformation (**?**), which would also suggest that $k_0$ must be chosen between $k_0 = 5 \cdot 10^5$ and $k_0 = 10^6$.

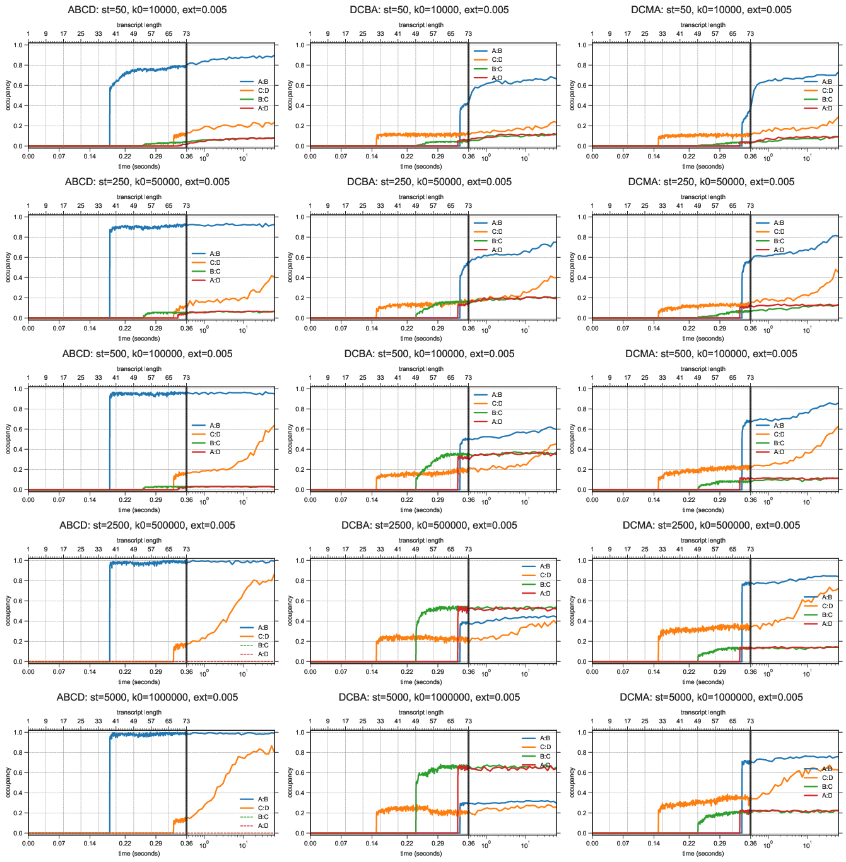


Figure 3: `Kinfold` simulations of the three molecules 'ABCD', 'DCBA', 'DCMA' designed in **?** with varying simulation time per nucleotide. The extension time per nucleotides is constant at **ext**=0.005 s/nt which corresonds to a transcription rate of 200 nt/s, which is unusually high, but was suggested in the original publication. $k_0$ is varied by two orders of magnitude: **k0**=$\{10^4, 5 \cdot 10^4, 10^5, 5 \cdot 10^5, 10^6\}$, where $10^5$ is the `DrTransformer` default parameter. The simulation time per nucleotide in arbitrary units **st** is provided for comparison to a different transcrption rate in Suppl. Fig. 5. The corrsponding `DrTransformer` simulations are shown in Suppl. Fig. 2. Experimental results for DCBA suggest 90% in B:C, A:D conformation and 10% in A:B, C:D conformation (**?**), which would suggest that $k_0$ must be chosen to be larger than $10^6$. Accordingly, experimental results for DCMA suggest 50% in B:C, A:D conformation and 50% in A:B, C:D conformation (**?**), which also suggests that $k_0$ must be chosen to be larger than $10^6$.

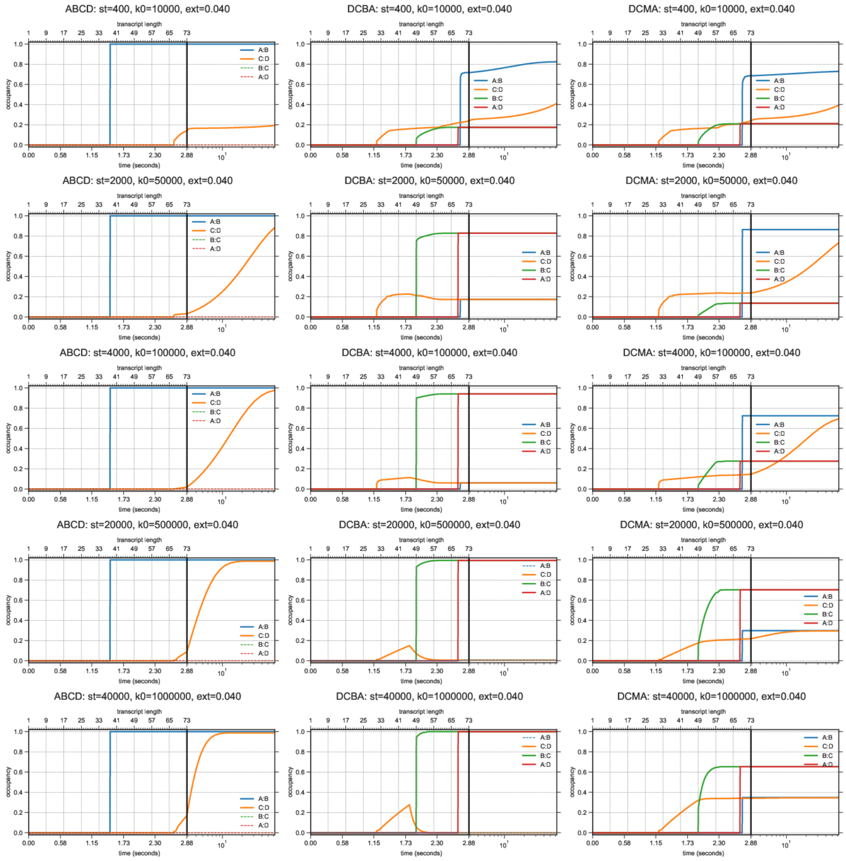


Figure 4: `DrTransformer` simulations of the three molecules 'ABCD', 'DCBA', 'DCMA' designed in **?** with varying simulation time per nucleotide. The extension time per nucleotides is constant at **ext**=0.040 s/nt which corresonds to a transcription rate of 25 nt/s (the `DrTransformer` default parameter). $k_0$ is varied by two orders of magnitude: **k0**=$\{10^4, 5 \cdot 10^4, 10^5, 5 \cdot 10^5, 10^6\}$, where $10^5$ is the `DrTransformer` default parameter. The simulation time per nucleotide in arbitrary units **st** is provided for comparison to a different transcrption rate in Suppl. Fig. 2. The corrsponding `Kinfold` simulations are shown in Suppl. Fig. 5. Experimental results for DCBA suggest 90% in B:C, A:D conformation and 10% in A:B, C:D conformation (**?**), which would suggest that `DrTransformer` default parameters ($k_0 = 10^5$ at 25 nt/s) give the best fit. Experimental results for DCMA suggest 50% in B:C, A:D conformation and 50% in A:B, C:D conformation (**?**), which would suggest that $k_0$ must be chosen between $k_0 = 10^5$ and $k_0 = 5 \cdot 10^5$, however, in practice is is difficult to get this 50/50 distribution for DCMA structures due to coarse graining effects.

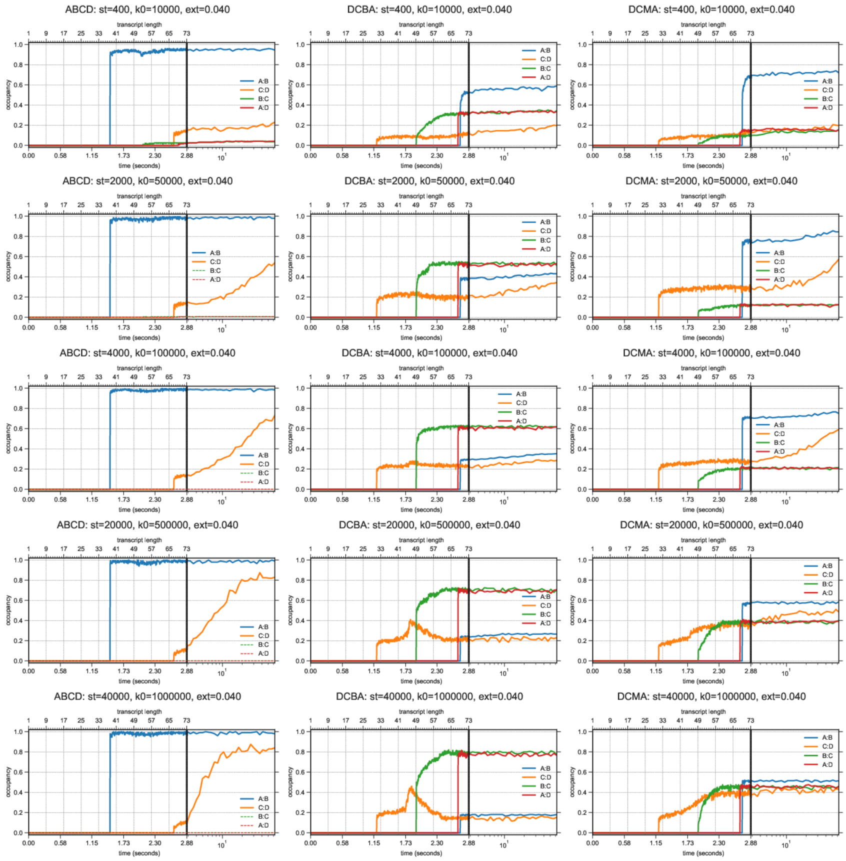


Figure 5: `Kinfold` simulations of the three molecules 'ABCD', 'DCBA', 'DCMA' designed in **?** with varying simulation time per nucleotide. The extension time per nucleotides is constant at **ext**=0.040 s/nt which corresonds to a transcription rate of 25 nt/s (the `DrTransformer` default parameter). $k_0$ is varied by two orders of magnitude: $\mathbf{k0}$=$\{10^4, 5\cdot10^4, 10^5, 5\cdot10^5, 10^6\}$, where $10^5$ is the `DrTransformer` default parameter. The simulation time per nucleotide in arbitrary units **st** is provided for comparison to a different transcrption rate in Suppl. Fig. 3. The corrsponding `DrTransformer` simulations are shown in Suppl. Fig. 4. Experimental results for DCBA suggest 90% in B:C, A:D conformation and 10% in A:B, C:D conformation (**?**), which would suggest that $k_0$ must be chosen to be larger than $10^6$. Experimental results for DCMA suggest 50% in B:C, A:D conformation and 50% in A:B, C:D conformation (**?**), which is well approximated by $k_0 = 10^6$.
